# Reproducibility in systems biology modelling

**DOI:** 10.1101/2020.08.07.239855

**Authors:** Krishna Tiwari, Sarubini Kananathan, Matthew G Roberts, Johannes P Meyer, Mohammad Umer Sharif Shohan, Ashley Xavier, Matthieu Maire, Ahmad Zyoud, Jinghao Men, Szeyi Ng, Tung V N Nguyen, Mihai Glont, Henning Hermjakob, Rahuman S. Malik-Sheriff

## Abstract

The reproducibility crisis has emerged as an important concern across many fields of science including life science, since many published results failed to reproduce. Systems biology modelling, which involves mathematical representation of biological processes to study complex system behaviour, was expected to be least affected by this crisis. While lack of reproducibility of experimental results and computational analysis could be a repercussion of several compounded factors, it was not fully understood why systems biology models with well-defined mathematical expressions fail to reproduce and how prevalent it is. Hence, we systematically attempted to reproduce **455** kinetic models of biological processes published in peer-reviewed research articles from **152** journals; which is collectively a work of about **1400** scientists from **49** countries. Our investigation revealed that about half (**49%**) of the models could not be reproduced using the information provided in the published manuscripts. With further effort, an additional 12% of the models could be reproduced either by empirical correction or support from authors. The other 37% remained non-reproducible models due to missing parameter values, missing initial concentration, inconsistent model structure, or a combination of these factors. Among the corresponding authors of the non-reproducible model we contacted, less than **30%** responded. Our analysis revealed that models published in journals across several fields of life science failed to reproduce, revealing a common problem in the peer-review process. Hence, we propose an 8-point reproducibility scorecard that can be used by authors, reviewers and journal editors to assess each model and address the reproducibility crisis.

## Introduction

Reproducibility of scientific results is a key determinant of true science and credibility. The reproducibility crisis across many fields of science has emerged as an important concern when results published by leading scientific journals failed to reproduce ^1–4^. In a survey of 1,576 scientists published in Nature ^5^, about 90% acknowledged this emerging crisis. The survey reported that over 70% scientists failed to reproduce others’ experiments and over 50% failed to reproduce their own results. Experimental results fail reproducibility tests due to several reasons including improper documentation of methodology, considering noise as a positive finding, unrecognized or incomplete experimental variables, data fabrication or bias, publishing premature or incomplete results ^6–9,5^ and inappropriate statistical analysis (e.g. “P-hacking”) ^10,11^. Computational biology research also faces this crisis, and the lack of reproducibility is compounded by several factors including changes in reference data and/or formats, software versions, missing essential codes or methodology etc. ^12–14^. Several suggestions have been published to improve reproducibility in computational biology ^12,15,16^ and bioinformatics ^17,18^.

Systems biology modelling involves mathematical representation of biological processes to investigate complex behaviour of the system which cannot be studied by looking at individual components ^19–21^. It was supposed that modelling will remain relatively untouched by this crisis, as the models are a specific set of computational codes representing well defined mathematical equations to perform reproducible simulations. However, mathematical models from a number of manuscripts do not reproduce the simulation results ^22–24^ described in the manuscript. While lack of reproducibility of experiment results and computational analysis could be a repercussion of several compounded factors, lack of reproducibility in the simulation of mathematical equations is typically due to inadvertent error or lack of information in the manuscript. It is critical to recognise the cause of models’ failure to surmount the reproducibility crisis. Moreover, it was also not clear how prevalent the reproducibility crisis is in the field of systems biology modelling.

Hence, to investigate the reproducibility crisis, we systematically analyzed mathematical models in conjunction with the curation process in BioModels repository ^25^. BioModels (https://www.ebi.ac.uk/biomodels/) is one of the largest public open source databases of quantitative mathematical models, where the models are manually curated and semantically enriched. In total we investigated 455 kinetic models of biological processes published in literature; which is collectively the work of about 1400 scientists from 49 countries. It is one of the largest studies performed to assess the status of reproducibility by attempting to independently reproduce results published in a number of research articles. We could reproduce about half of the models (51%), whereas the other half could not be reproduced using the information provided in the manuscript. We managed to reproduce a further 9% of the models with empirical trial and error approach and an additional 3% with the help of authors who responded to our email requests. Our in-depth investigation has revealed the major reasons why models fail to reproduce the published simulation results. We discuss these reasons and further propose a reproducibility scorecard for modellers, reviewers and journal editors to assess models and address the reproducibility crisis.

## Methodology

To investigate the reproducibility in the systems biology models, we systematically analyzed 455 ordinary differential equation (ODE) models from published research articles. This exercise was done in conjunction with the curation of models in BioModels. The manual curation of models in BioModels involves a two-step process (1) encoding models in standard formats and reproducing the simulation figures in the reference manuscript and (2) semantic enrichment of the model and its components ^25^. The reproducibility assessment was done along with the first step of the model curation. Semantic enrichment of the model in BioModels was done following MIRIAM guidelines ^26^ and it involved annotation of model entities (species, reactions, parameters, events, etc) with cross-references to controlled vocabularies such as GO ^27,28^, ChEBI ^29^, Mathematical Modelling Ontology ^30^, Systems Biology Ontology ^31^, Brenda Tissue Ontology ^32^ and Experimental Factor Ontology ^33^, as well as data resources such as UniProt ^34^, Ensembl ^35^, NCBI Taxonomy ^36^, Reactome ^37^, etc. The models for curation are selected either from those submitted to BioModels by modellers or from the published articles contingent on the interest of the curators and collaborators. Our collection covered models of a wide range of biological processes, originating from articles published in 152 life science journals.

### Reproducibility assessment

The following steps were employed in our study to assess the reproducibility:

1. The model manuscript was carefully read and the model equations were encoded in the standard SBML ^38^ format. When the models were previously submitted in SBML format, the equations, values of parameters and initial concentration, perturbation events, etc. were cross-verified with the reference manuscript.
2. The simulation of SBML model files was performed predominantly using COPASI ^39^. When COPASI was used in the original manuscript, other simulation software such as simbiology toolbox (MATLAB) ^40^, libSBMLsim ^41^, Mathematica ^42^ were used to perform simulations.
3. The model was considered as reproducible when it reproduced at least one of the main simulation figures in the associated research article using a software different from the one used in the original manuscript. The reproduced simulation figure, such as time course plot with and without perturbation, phase-plane plot, etc., should precisely match the original figure, any minor deviation was still considered acceptable if it did not affect the scientific conclusion of the study. The models which could be directly reproduced with the description in the manuscript were labelled as ‘Directly Reproducible’.
4. When the model failed to reproduce the simulation with the mathematical equation and the parameter values provided in the research article, we resorted to an empirical trial and error approach to correct the model based on curator expertise. For example, any terms missing in the equations but described in the manuscript were added to correct the model; any potential typos such as misplacement of decimal points in the parameter values were corrected. The models that were reproduced after such corrections were labelled ‘Reproduced with manual corrections.
5. The authors of the failed, yet potentially salvageable models were contacted when possible and their responses were recorded. The models that were reproducible with the corrections provided by them were labelled as ‘Reproduced with author support’.
6. Nevertheless, when the model still could not be reproduced, they were labelled as ‘Non-reproducible’ and the likely reasons were recorded. The plausible reasons for non-reproducibility include (a) inconsistency in model structure i.e. any error in the model equation (b) missing parameter values (c) missing initial concentration and (d) unknown reason.
7. SBML representation of all the analysed models were submitted to BioModels ^25^. The reproducible ones were labelled as curated models.

## Results

### About half of the published models were not directly reproducible

In total 455 kinetic models from peer-reviewed and published scientific publications were subjected to reproducibility tests and extensively analysed to unveil the fundamental cause behind systems biology model’s failure to reproduce the simulation figures in the associated manuscripts. We selected kinetic model manuscripts published from 1980 to 2020 (Sup Fig 1). Among these models, the mathematical equations of 389 models were manually encoded in the standard SBML format using COPASI ^39^ from the original manuscript. The remaining 66 models were those submitted to BioModels in SBML format by the authors and hence they were carefully cross-checked to ensure whether the mathematical equations, initial conditions and parameters were accurately represented. The SBML representation of 233 out of 455 models (51%) directly reproduced simulation results in the original manuscript (Figure 1a). About 49% of the published models were not reproducible either due to incorrect or missing information in the manuscript. This high proportion is unexpected and exposed a serious issue within the field. Even among the 66 models submitted to BioModels in SBML standard format, only 37 could be reproduced directly.

**Figure 1:**
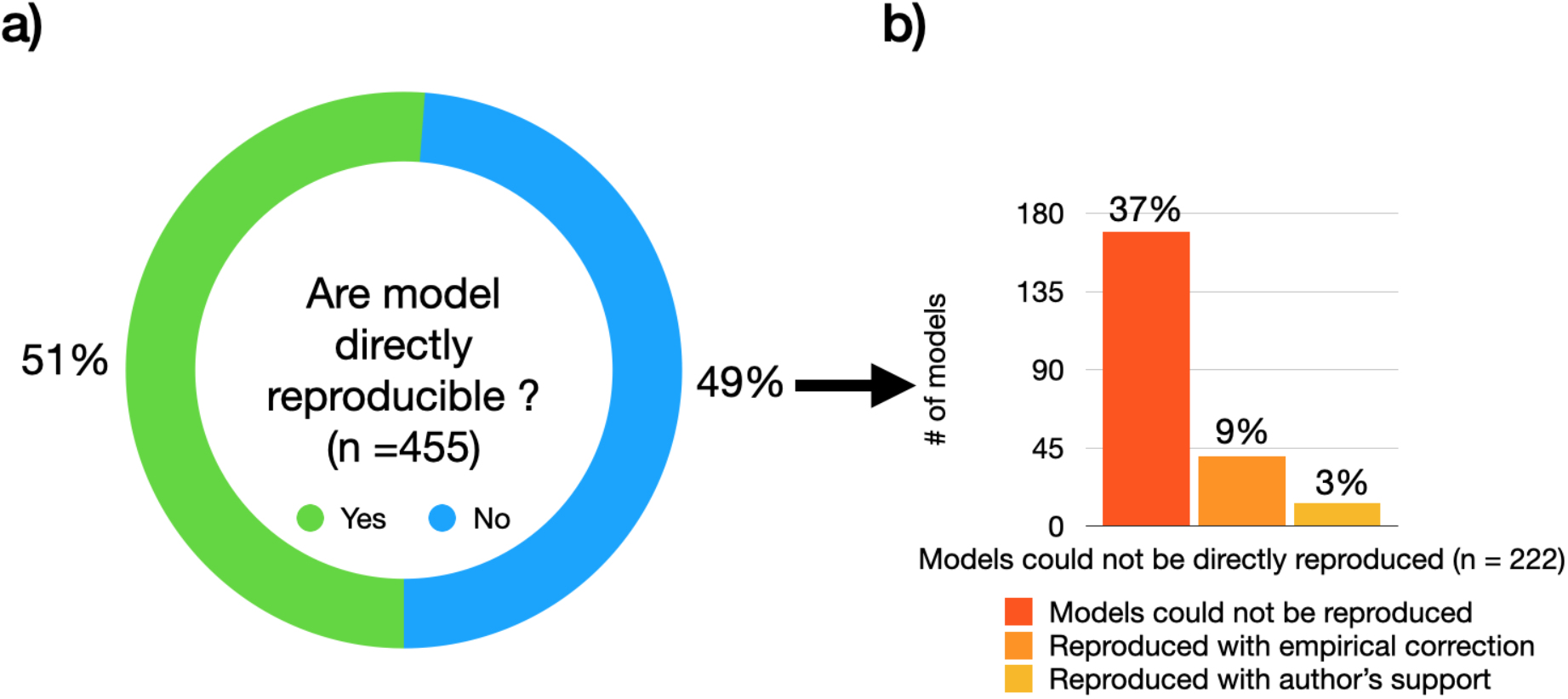
Reproducibility of systems biology models. (a) About half of the published systems biology models could not be directly reproduced. (b) About 12% of the models could be reproduced with empirical corrections or author support.

### About 12% of the models could be reproduced with further effort

About 12% of all models could be rescued with further efforts involving either a careful empirical trial and error approach or author support (Figure 1b). 40 models (9%) were successfully reproduced with manual empirical correction of the inaccurate reporting in the manuscript. Some of the common errors that could be identified and manually corrected were (a) error in the sign of the terms in the mathematical equations, e.g. a negative sign for a production term in the equation or vice versa, could be easily identified by expert curation and corrected (b) missing terms in the model equations - e.g. missing one of the production or depletion terms in the ODE definition (c) typos in the parameter values e.g. mistakes in the decimal points, a value of 0.01 was reported in place of 0.001 (d) missing values e.g. some missing initial concentrations of model entities could be inferred from the initial time point in the simulation plots (e) error in the units of initial concentration and parameter values e.g. nM/l was misrepresented as uM/l.

It was not always possible to estimate the missing concentration, parameter or the mathematical expression. Hence, we contacted corresponding authors to request the missing information or seek clarification. This was not always feasible, for example due to authors’ change of institutions, change of field, leaving academia, and death. In total, we attempted to contact corresponding authors of 90 models, among which less than a third (27) responded (Figure 2). About half of the models of authors who responded (13 models, or 3% of the total number of models) were subsequently reproduced. These were mostly models published in the last 5 years (Sup Fig 1). Surprisingly about 70% of the authors we contacted did not respond to the request to provide information to reproduce their models (Figure 2).

**Figure 2:**
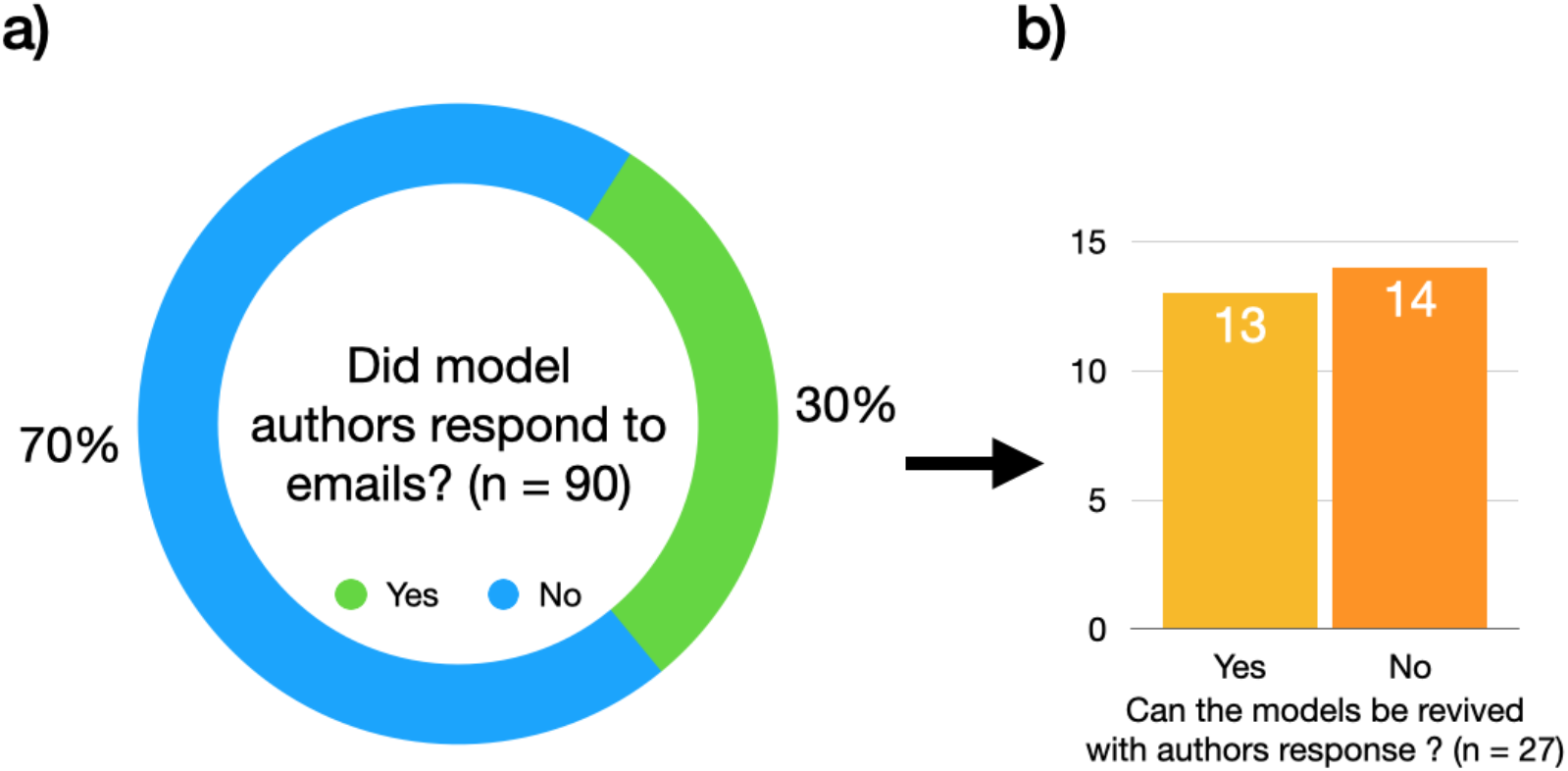
Rescuing reproducibility with author support. (a) A majority of authors of non-reproducible models did not respond to emails. (b) About half of the models of the authors who responded could eventually be reproduced.

### The major reasons why models failed to reproduce

About 37% of the 455 models could not be rescued even with further efforts. The major reasons why these models failed to reproduce include missing parameters values as the main reason, followed by missing initial concentration and inconsistency in model structure, or a combination of aforementioned causes (Figure 3). Yet, in a large proportion of the non-reproducible models, the reason for failure was unclear. The reference manuscripts of those models might have probably reported incorrect parameter values, initial concentrations or model equations or a combination of these three factors. Some research articles report parameter values in the form of plots, and it was not straightforward to extract those values to reproduce the simulations. Insufficient and incorrect reporting of the model content were the main reasons why models failed to reproduce. These factors are commonly overlooked in the peer-review process and hence are reflected across several journals where systems biology models are published (Figure 4).

**Figure 3:**
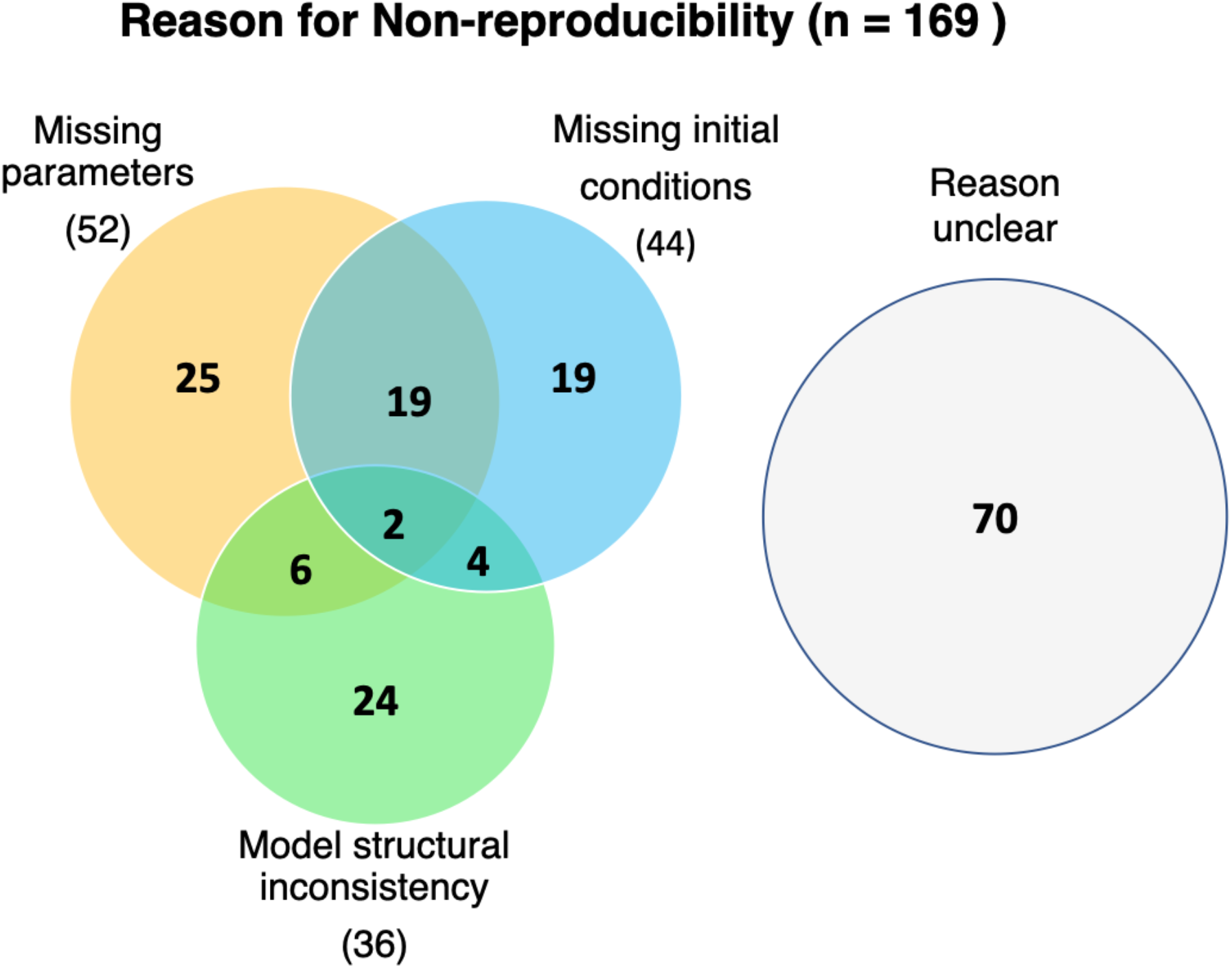
Common reasons why models could not be reproduced include missing parameter values, initial conditions and inconsistency in model structure.

**Figure 4:**
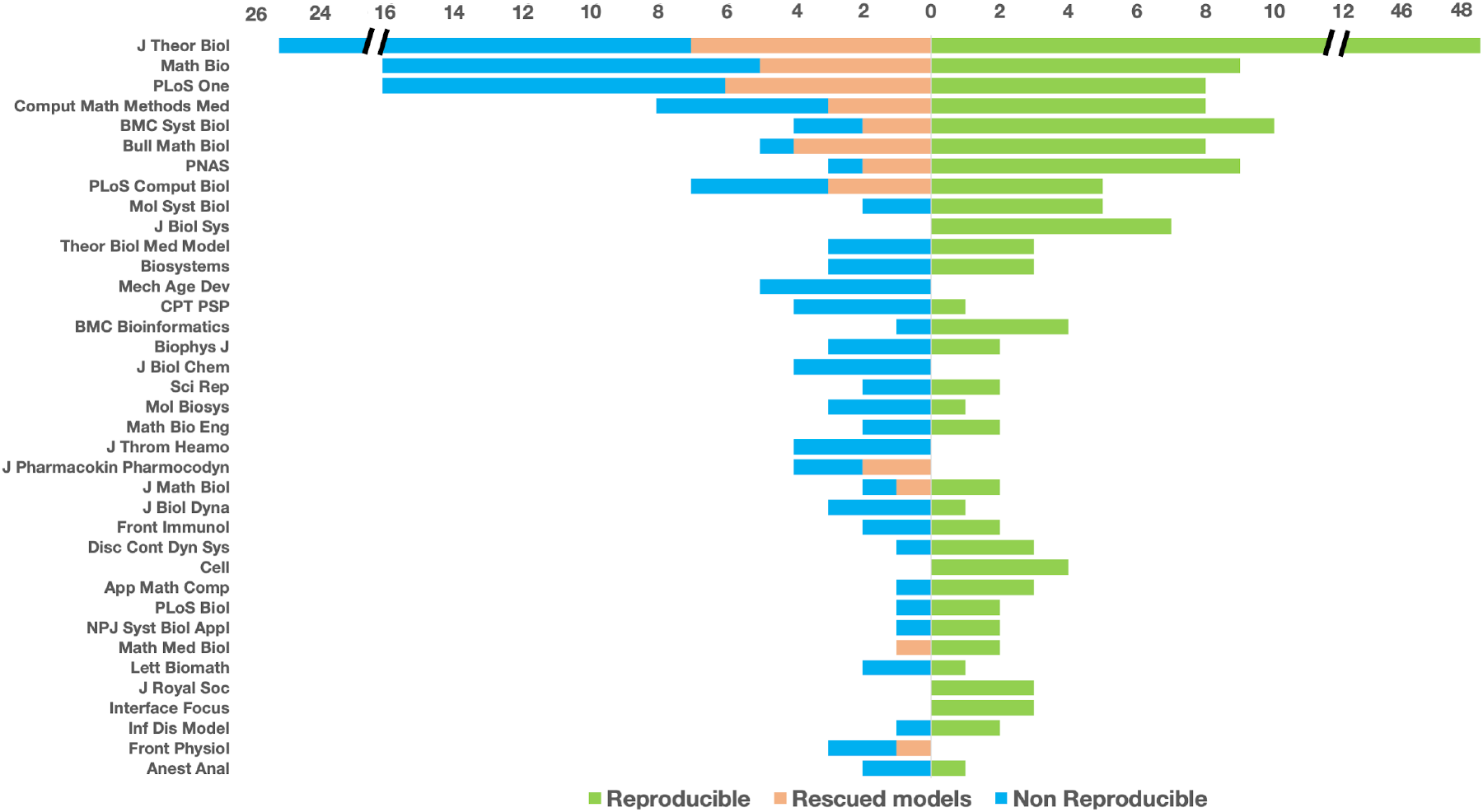
Distribution of reproducible, rescued and non-reproducible models across various journals with more than two models in our study.

## Discussion

Reproducibility is the essence of science and any scientific work that cannot be reproduced independently using the same resources is deemed untrustworthy ^22,43^. The ability to reproduce a biological experiment is often limited by several confounding factors and is not unheard of. Similarly, reproducible bioinformatics analysis is affected by regular updates, releases in data and software, etc.^18^. Systems biology models are least expected to be affected by the lack of reproducibility, as they are merely a set of mathematical functions that are straight forward to simulate. In contrast, our analysis of 455 ODE models has exposed that about half of these models cannot be reproduced using the information provided in the manuscript. Although the lack of reproducibility has been discussed within the community ^17,12,22^, it was never expected to be this adverse. Our study has revealed that the inability to reproduce models is widespread in research articles published across several journals in the field with the exception of a few (Figure 4), and it is imperative to revisit the peer-review process of mathematical studies as previously suggested ^23^.

Reproducibility, replicability, repeatability are the terminologies often confused and are defined differently in experimental ^43,5^ and computational research ^44^. In the context of systems biology modelling, the refined definition of replicability (also referred as repeatability) is the ability to use the same code provided with the manuscript in the same software to reproduce the simulation results; whereas reproducibility is the ability to build the code *de novo* and/or ensure the mathematical expressions are correctly represented and reproduce the simulation results in a software different from the one originally used. The focus of this work was to assess the reproducibility of the published systems biology models and hence the latter was chosen as the criterion.

Even in the models that are reproduced, one of the challenges we faced is unambiguous inference of the model entities and their values. When a variable or model entity name is different in the main manuscript description, mathematical expression, and code, it becomes challenging to match them to reproduce the simulation. For example, “alpha” in the model description and/or equation in the manuscript and the code may refer to completely different entities. We have overcome this challenge by carefully reading the reference manuscript. We strongly recommend making the code or the model file as self-contained as possible with proper annotation of the model entities. Model codes written in programming languages such as Matlab, python, C, R, etc. are often helpful to reproduce the model. Nevertheless, not all such codes are easily comprehensible, specifically when they are not well commented. Although the systems modelling community is split, a notable fraction of the modellers use COMBINE ^45^ community standard formats such as SBML, SED-ML ^46^, COMBINE Archive to encode their models. These standard formats provide a consistent framework to encode and annotate models, making them both human and machine readable. The strong community support for standard formats such as SBML makes it highly interoperable with about 280 supporting software tools for model construction, simulation, visualization and processing the semantic layer. We highly recommend using standard formats to encode and disseminate mathematical models as these greatly enhance the ability to comprehend and reproduce the models.

The most common approaches in systems biology modelling include kinetic, constraint-based, logic and agent-based modelling ^21^. In this work, we specifically focused on ODE models, one of the types of kinetic models, to analyse why they cannot be reproduced. Half of these relatively simple deterministic kinetic models could not be reproduced. Other types of kinetic models include delay differential equations and partial differential equations; they are likely to be affected either to the same extent or even more by the reproducibility crisis as they are relatively more complex than ODE models. In the case of constraint-based modelling, flux values resulting from flux balance analysis are commonly reported in manuscripts and these values are not unique solutions and cannot be directly reproduced. The community tool MEMOTE ^47^ was developed primarily to quality control constraint based models. We are currently collaborating with the constraint-based modellers to develop tools and procedures to test reproducibility of these models. Similarly, we have engaged with the logic modelling community to develop guidelines for curation and annotation of logic models (CALM) ^48^.

Leveraging on the lessons learned from attempting to reproduce 455 models as well as our interaction with modelling communities, we have developed a reproducibility scorecard to enhance the ability to reproduce systems biology models (Box 1). The scorecard consists of a list of items that would allow another modeller to reproduce the simulation results of a model with a reasonable effort. We recommend authors, reviewers and editors of the journals to assess each systems biology model in the research article using our proposed scorecard. The scorecard consists of 8 questions with a unit score for each ‘yes’ as an answer. All the 8 questions may not always be applicable and hence, in the scale of 8, we strongly advocate that a model get a score of 4 at the least.

A clear and complete description of the mathematical model, relevant parameter values and simulation conditions are vital to reproduce the model and are addressed in the first three questions. In case of large models, it may not be possible to enumerate the aforementioned information in the manuscript, hence it is acceptable to share the code publicly either via supplementary material, git-hub, or a model repository. Following the Findable, Accessible, Interoperable and Reusable (FAIR) ^49^ principles, we have recommended submission of the model code in standard formats into the relevant model repositories in question 5 and 6 respectively. However, question 4 is not redundant, as it is used to cover those modellers who use non-standard formats, for example as collection of programming scripts and share them via git-hub, unstructured repositories, or similar. A model shared in standard format such as SBML, CellML ^50^ gets an additional score in our scorecard, as the interoperability of the model file will provide the possibility to test and reproduce it using several supporting software tools.

#### Box1 Reproducibility score card

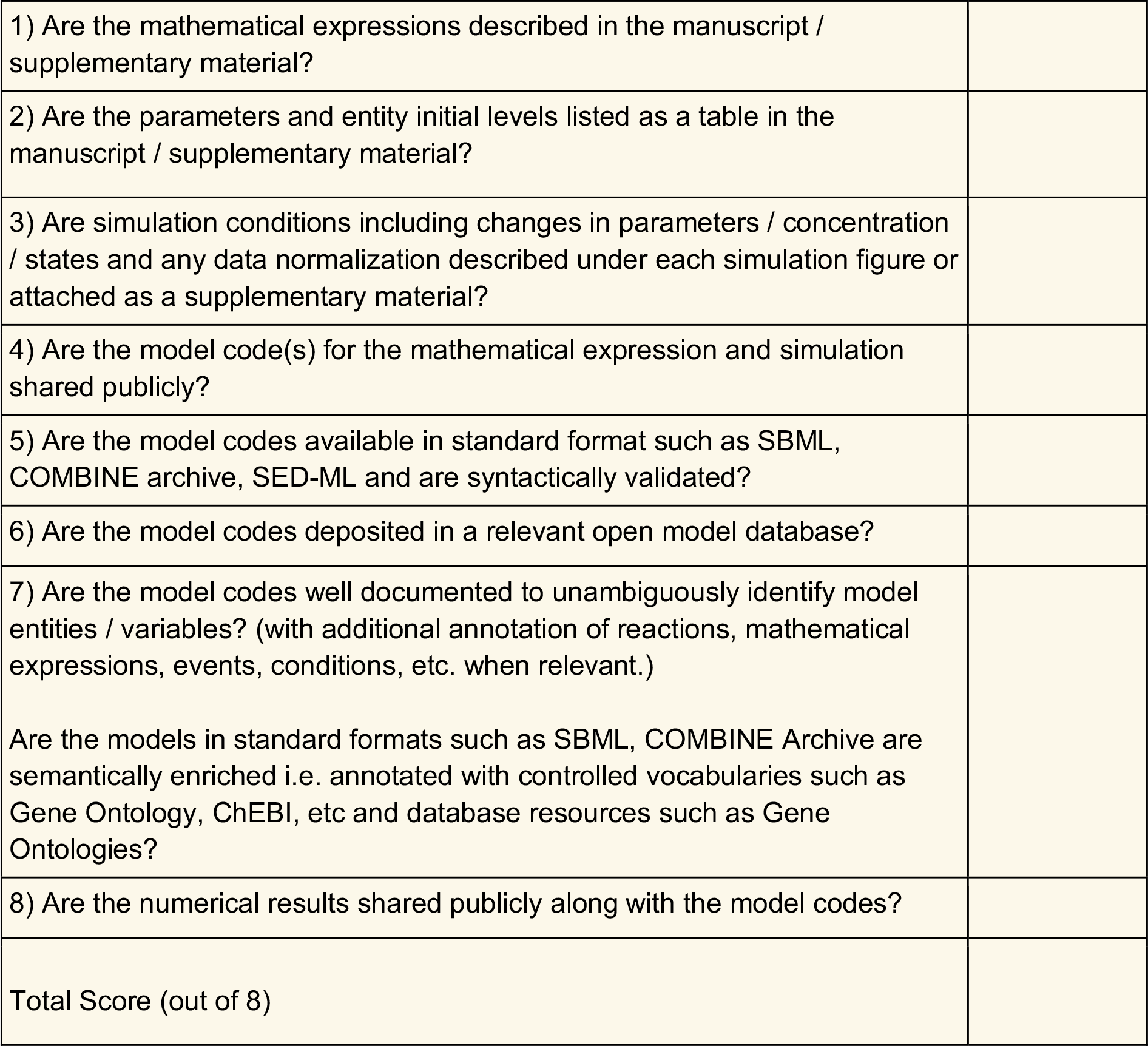

Similarly, deposition of models to open repositories including, but not limited to BioModels, Physiome ^50^ or JWSOnline ^51^ will get an additional score as they promote FAIR sharing. The advantage of submitting models to open model repositories include provision of (1) a sophisticated search engine to make models findable, (2) a version-controlled storage system to make the models readily accessible, (3) support for interoperable standard formats and (4) curation and annotation services to promote reusability. BioModels, being the largest repository of open, curated, well annotated, and findable models, contributes significantly to reproducibility ^22^. Several journals recommend to submit their model to BioModels ^52^. Similar to the curation service in BioModels, JWSOnline and the Physiome repository through the Centre for Reproducible BioMedical modelling ^12^ provide expert curation services to validate published reports.

Unambiguous identification of the model entities is critical to reproduce the results, hence semantic enrichment or proper documentation of the code brings added value. Even an accurately defined model cannot be reproduced if the data normalization and the simulation conditions which include changes to specific parameters or concentration of model entity are not clearly described. Hence, we recommend provision of this information under each simulation figure. Alternatively, the modellers can submit SED-ML files, a COMBINE standard for description of simulation experiments. Although it is possible to use the simulation figures in the manuscript as reference to test the reproducibility of the models in many cases, it is desirable to provide the numerical output of the simulations to verify the model reproducibility.

Model curation is a time intensive task; on average it took about a week to carefully encode and thoroughly investigate the reproducibility of a model; in some cases, it took less than two days and in some cases over two weeks. Our criterion for a reproducible model was that it should reproduce at least one figure from the original article, which we consider to be a reasonable compromise between reliably assessing reproducibility, and the huge additional effort that would be required to ensure reproducibility of all (relevant) figures.

Our study was not intended to call-out non-reproducible models, but rather to highlight the current status of and reasons behind the lack of reproducibility. Thereby, we intend to raise community awareness among the researchers who use systems biology models and provide a potential solution to address the reproducibility crisis. We provide the “curated” section of BioModels as a source of reliable, verified reproducible models, while still keeping the non-reproducible models accessible. The versioning system in BioModels will allow authors to improve their models, while keeping the original version accessible as a part of the public record. Currently, we just keep the non-reproducible models in the “non-curated” part of BioModels, which also contains models from direct submissions which are still awaiting curation, as well as models in representations for which we currently don’t provide detailed curation. We do not explicitly label non-reproducible models because, on the one hand, there is a chance that the failure to reproduce the model is due to curator error, and on the other hand we don’t want to discourage authors from making their models accessible through a public repository. However, we are aware that the lack of explicit labelling of a non-reproducible model might cause others to try the same again. We are open to community suggestions for a more transparent labelling of models.

Lack of reproducibility has an adverse effect on the reliability of the scientific results. An additional consequence is the loss of the time and resources spent by modellers across the globe attempting to reproduce a previously published model. Availability of model code can reduce the time needed to reproduce the model. Nevertheless, submission of model codes in standard format to open repositories is not alone sufficient to reproduce. Among the models we investigated were 66 SBML models submitted by modellers to BioModels, yet nearly half of them (29) could not be reproduced directly either. Hence, we formulated the 8-points reproducibility scorecard with a 4-point cut-off. BioModels will actively seek to curate models published with our proposed score card. By adapting our scorecard in the peer-review process, complemented by the curation services provided by model repositories and reproducibility centres, we believe that the reproducibility crisis can be significantly addressed. It can be achieved as a community, where authors, reviewers, journal editors embrace reproducibility more proactively than before.

