## Supplementary Figure 1 for "Reproducibility in systems biology modelling"

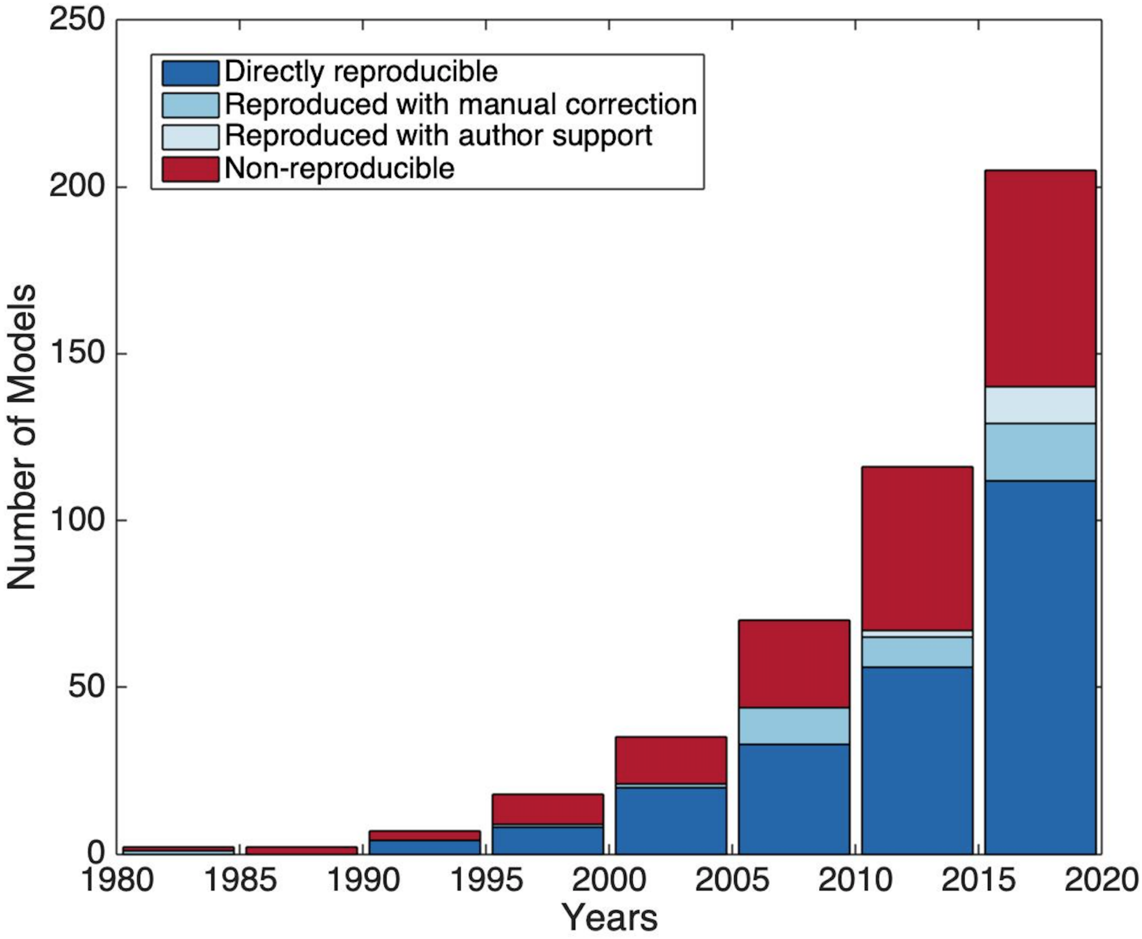


Supplementary Figure 1: The year of publication of the models analysed in this study.
